# High-resolution phenotypic screen in zebrafish identifies novel regulators of CNS axon diameter growth

**DOI:** 10.1101/2025.04.29.651302

**Authors:** Maria A Eichel-Vogel, Daniel Soong, Melody N Sequeira, Katy LH Marshall-Phelps, Jenea M Bin, David A Lyons

## Abstract

Axon diameter varies up to 100-fold between distinct neurons in the central nervous system with larger axons exhibiting proportionally faster conduction velocity. Axon diameter influences myelination, can be dynamically regulated, which might help fine-tune neural circuit function, and is altered in several diseases. Despite its importance, mechanisms regulating axon diameter remain poorly understood. This gap in understanding is due in part to the limitations of fixed tissue analyses, such as electron microscopy, which cannot be scaled up to execute discovery screens. To address this, we developed a high-resolution, high-content imaging-based in vivo platform to identify pharmacological modulators of axon diameter in zebrafish. We focused on the Mauthner neuron, whose axon diameter growth can be monitored during development. To facilitate our high-content chemical screen, we developed an automated high-resolution imaging and image analysis pipeline to assess changes in Mauthner axon diameter in transgenic reporter animals. We screened 880 compounds and identified 33 that altered Mauthner axon diameter. Validating this discovery pipeline, we confirmed that compounds that affect beta-2 adrenoceptor and dopamine signaling increase axon diameter in separate follow-up studies. This represents the first discovery screen for axon diameter regulators, providing novel entry points to study the biology of axon diameter regulation.

## Introduction

Across the vertebrate nervous system the diameter of axons can vary by over 100-fold, correlating with a roughly corresponding 100-fold variation in action potential conduction speed, with larger axons propagating impulses faster. The range in diameter equates to ∼10,000-fold difference in area occupied by distinct axons^1–4^. In addition to differences between axons, diameter can vary along individual axons, including between branches, and over time. Indeed, evidence suggests that axonal diameter may mediate the precise conduction properties of specific circuits and that it can be dynamically regulated via cell-cell interactions and neuronal activity^5–13^. Furthermore, axon diameter is altered in a range of diseases and injuries, the cause and consequences of which remain to be determined^14–21^.

Driven largely by static electron microscopy-based studies, a relationship between myelination and axon diameter has been evident for over a hundred years^22–28^, and between the axonal cytoskeleton and axon diameter for over fifty years^29–32^. Recently, innovations in super-resolution light microscopy have led to the identification of additional features of the axonal cytoskeleton that influence axon diameter^33–34^, but widespread investigations of axon diameter biology have not yet been undertaken. This is, in part, due to the relatively low throughput of the very high-resolution approaches typically required to assess diameter in intact tissue. As such, there have not yet been any scalable discovery-based screens applied to axon diameter in model systems such as *C elegans*, *Drosophila* or zebrafish, despite the fact that such screens have taught us much about other features of axon biology ^35–42^.

Recently, we established zebrafish as a model to study axon diameter and used transgenic reporters to observe axon diameter growth in vivo over time^43^. Following a genetic screen for new genes required for myelinated axon formation, we identified the nuclear transport receptor ipo13 as being essential for the growth in diameter of very large axons. This capacity motivated us to develop a discovery platform that aimed to identify novel regulators of axon diameter biology, a feature of neuronal morphology and nervous system function that has been relatively understudied to date. To perform a screen for axon diameter, we reasoned that it would be critical to 1) use a system that allows us to investigate even subtle changes to axon diameter as precisely as possible; 2) track and resolve individual axons in vivo; 3) reliably and robustly image and quantify even subtle changes to diameter and 4) automate this process to increase scalability for screening hundreds of compounds. Our previous studies had shown that the Mauthner axon grows by several microns in diameter in the days following its outgrowth^43^. Here we describe the development of an automated imaging and machine learning-based analysis pipeline to assess changes to Mauthner axon diameter in the zebrafish, the execution of a large chemical-compound based discovery screen, and validation of hits. We also provide our findings as a resource for the community to accelerate investigation into the biology of axon diameter.

## Results

### Automated high-resolution imaging of axon diameter in vivo using zebrafish larvae

To investigate the molecular regulation of axon diameter in the CNS, we chose the Mauthner neurons and their axons as a model. Mauthner neurons are a pair of reticulospinal CNS neurons critical for mediating escape responses that project long large diameter axons from the brain to the spinal cord^44–46^. Our previous assessments of the Mauthner axon employed super-resolution confocal live imaging of fluorescent reporters and electron microscopy and indicated the axon grows from ∼1µm to ∼5µm in diameter between 2- and 5-days post fertilization. However, neither super-resolution live imaging nor electron microscopy allow the experimental throughput needed for a discovery project. Furthermore, our previous measurements of axon diameter were carried out either manually or using a semi-automated analysis pipeline, also unsuitable to scalable analysis of changes to a sub-cellular parameter, not detectable by eye. Therefore, to implement a scalable discovery screen, we reasoned that we needed to automate both the imaging and subsequent analysis of Mauthner axon diameter in a manner that balanced imaging capable of detecting subtle changes with the throughput required to execute a screen.

We previously employed automated live imaging of zebrafish expressing fluorescent reporters to screen for compounds that influenced myelinating glial cell number^47^. To do so we had employed a spinning disk confocal microscope (SDCM) for image acquisition, coupled to the vertebrate automated screening technology (VAST) platform for the automated delivery of zebrafish larvae from the wells of a 96-well plate to the microscope for imaging. Therefore, we wanted to determine whether this high-throughput platform could acquire images at a sufficient resolution to detect changes in the diameter of the Mauthner axon.

The VAST platform delivers zebrafish to a thin-walled glass capillary (600-750µm bore), in which they are stabilized and automatically oriented to visualize the animal from a specific orientation. The system then hands control to the SDCM to acquire images of the animal within the glass capillary. As previously, we used the transgenic reporter Tg(hspGFF62A:Gal4); Tg(UAS:GFP)^41,48^ to image growth in the diameter of the Mauthner axon. Due to fact that Mauthner axons are located in the ventral spinal cord close to midline axis of the animal, adjacent to the central canal, we needed to use long-working distance objective to image through both the tissue overlying the axons and the glass capillary, thereby limiting the objectives to relatively low-magnification water dipping lenses. We first tested a 10x 0.5NA WD=3.7mm objective and acquired 6 tiled z-stacks to image the whole animal, as per our previous myelinating glial cell screen, but found that this did not provide the resolution required to satisfactorily image the Mauthner axon (Figure 1A and A‘). In addition, delivery of fish to the capillary and imaging those 6 z-stacks took 4.45 minutes for one individual fish. With these settings a 96-well plate with 1 fish/well would take over 7 hours to image. To achieve higher resolution and minimize image acquisition time, we chose to change our imaging parameters: 1) we acquired images using a 20X 1.0NA WD=2.3mm objective and 2) we collected just one confocal z-stack in a single region halfway down the spinal cord, where we have previously assessed axon diameter. To ensure that we imaged the same region every time, we acquired the z-stack at 1900µm from the tip of the nose (Figure 1B and see Methods). Compared to imaging the whole animal, this approach allowed both delivery of the animal from the well to the capillary, orientation, and imaging to be carried out in ∼ 2 minutes 30 seconds. Table 1 summarizes the different resolution and imaging times based on one fish/well.

**Figure 1:**
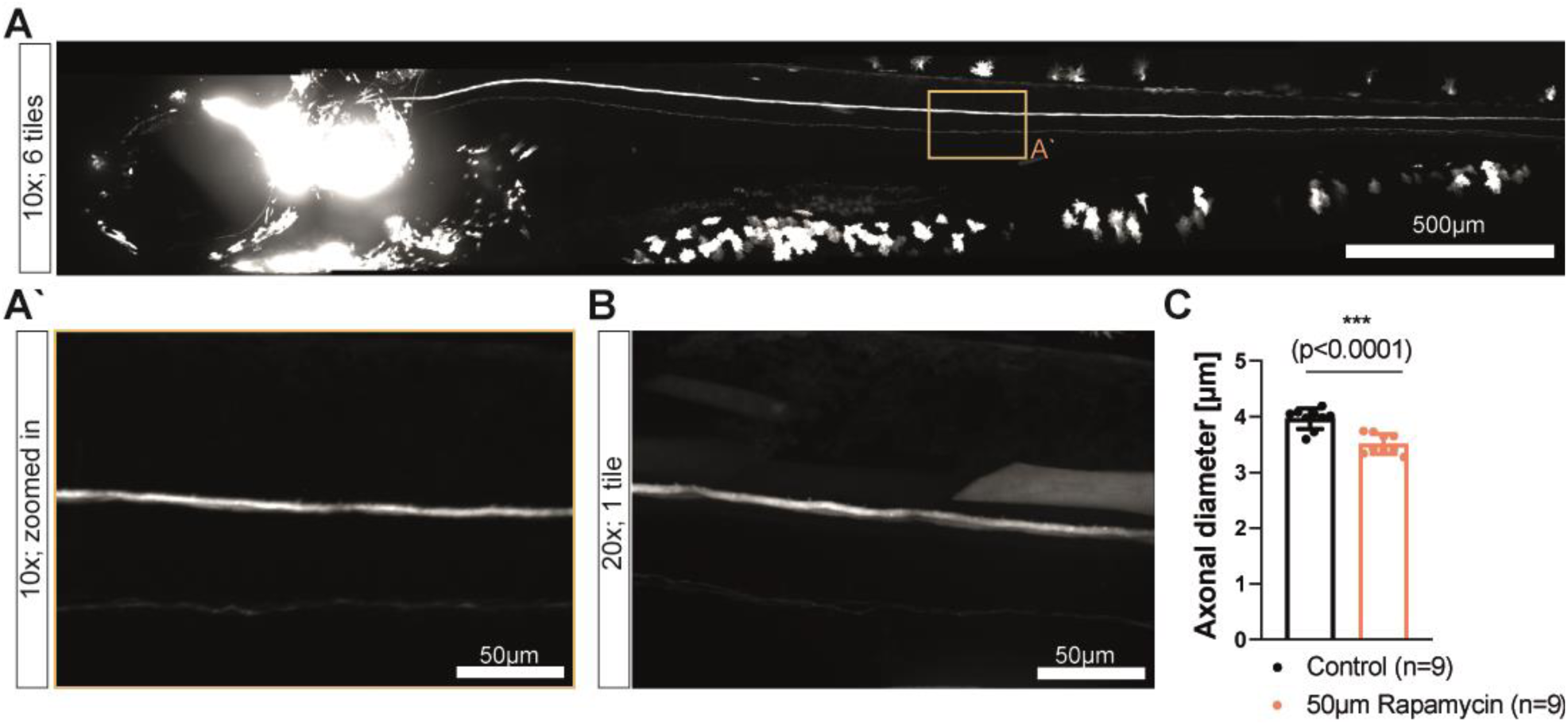
Axon diameter changes can be visualized and assessed using an automated screening system and analysis pipeline. A) Maximum intensity projection of images of Tg(Mauthner:EGFP) zebrafish following tiled merging of 6 z-stacks. Z-stacks were acquired from lateral view with a 10x/0.5NA M27 75mm objective using VAST-SDCM system. Scale bar = 500µm). À) Zoom in from the maximum intensity projection of Tg(Mauthner:EGFP) at 10x indicates that resolution does not meet anticipated standard for measuring axon diameter differences. B) Maximum intensity projection of 1 z-stack at approximately somite 15-16 of Tg(Mauthner:EGFP) fish imaged with a 20x 1NA (1×1 binning) objective shows improved subcellular resolution compared to 10x. C) Script based semi-automatic quantifications of Mauthner axon diameter show that we can measure axon diameter changes using the imaging parameters in B. When applying the mTor inhibitor Rapamycin we detect a significant decrease in Mauthner axon diameter (n=9, ***p<0.0001).

**Table 1.** Imaging resolution and times for 1×1 binning levels using either 10x or 20x objective and imaging Tg(hspGFF62A:Gal4); Tg(UAS:GFP) reporter line at 300ms exposure.

|  | **10x** | **20x** |
| --- | --- | --- |
| Pixel size (2x2 bin) | 0.556µm | 0.286µm |
| Exposure | 300ms | 300ms |
| Fish loading time (s) 1 fish/well | 50 | 50 |
| Fish loading time (s) 3 fish/well | 30 | 30 |
| Load and image complete fish (min) | 4.45 (6 tiles) | N/A |
| Load and image 1 tile z-stack (min) | 2.30 | 2.30 |

Next, we wanted to test whether images acquired in this way allowed us to detect experimentally induced changes in axon diameter, and to determine how many animals would be needed to detect statistically significant changes to diameter upon manipulation. To detect changes to the growth of axons in diameter and not earlier stages of neuronal or axonal maturation, we wanted to carry out manipulations after the Mauthner axon had completed its outgrowth and during active stages of diameter growth (3-5dpf). To achieve interventions during this time window, we selected a chemical screening approach of well-defined compounds as our primary strategy for identifying potential mediators of axon diameter growth since compounds can be easily applied by immersion at desired developmental timepoints. To validate this premise ahead of carrying out a screen, we assessed whether we could detect changes in axon diameter induced by manipulating one of the few signaling pathways previously implicated in influencing the diameter of myelinated axons. Due to its global regulation of cell growth, the mTor pathway has been shown to affect axon diameter, including in a mTor mutant zebrafish (data presented but not analysed in a study of myelination by Kearns et al., 2015^49^). To detect potential changes to axon diameter upon chemical manipulation of mTor we applied the inhibitor Rapamycin at 50µM from 3-5dpf, during active diameter growth. Using our candidate imaging parameters at VAST-SDCM and our previously described semi-automated script to quantify diameters^41^, we were indeed able to detect a decrease in Mauthner axon diameter of around 10% upon Rapamycin treatment. This was statistically significant when using 9 animals, indicating the sample size that we would likely need to detect such subtle changes to diameter in the context of a screen (Figure 1C).

### Creating an automated image analysis pipeline to measure axon diameter changes at subcellular resolution

Having established that the VAST-SDCM system can reliably detect a 10% change in axon diameter at 5dpf, we next wanted to develop an image analysis pipeline to fully automate the quantification of axon diameter. The transgenic line Tg(hspGFF62A:Gal4); Tg(UAS:GFP) that we used to label the Mauthner axon also labels neurons in the posterior lateral line nerve and, on occasion exhibits ectopic fluorescence in muscles and other skin cells, as is common for transgenic zebrafish with Gal4:UAS-driven expression (Figure 1A; 2A). Variation in such ectopic fluorescence expression, as well as autofluorescence, skin pigmentation, and biological batch/clutch variations, all lead to an extensive range of inter-image variability and quality. Valid expression of GFP in axons in one larva can sometimes be dimmer than undesired artefacts in another larva. Furthermore, although the VAST system orients the zebrafish in a lateral position with great fidelity, it is not possible to ensure that the midline of the animal runs parallel to the midline of the capillary tubing. Therefore, the trajectory of the Mauthner axon is not always constrained to a predictable number of z-sections in acquired stacks. For these reasons, it was not possible to simply convert 3D z-stacks into 2D maximum intensity projections for threshold-segmented analysis. Instead, we employed machine learning to segment objects of interest in manner that is more agnostic to intensity or position (Figure 2).

**Figure 2:**
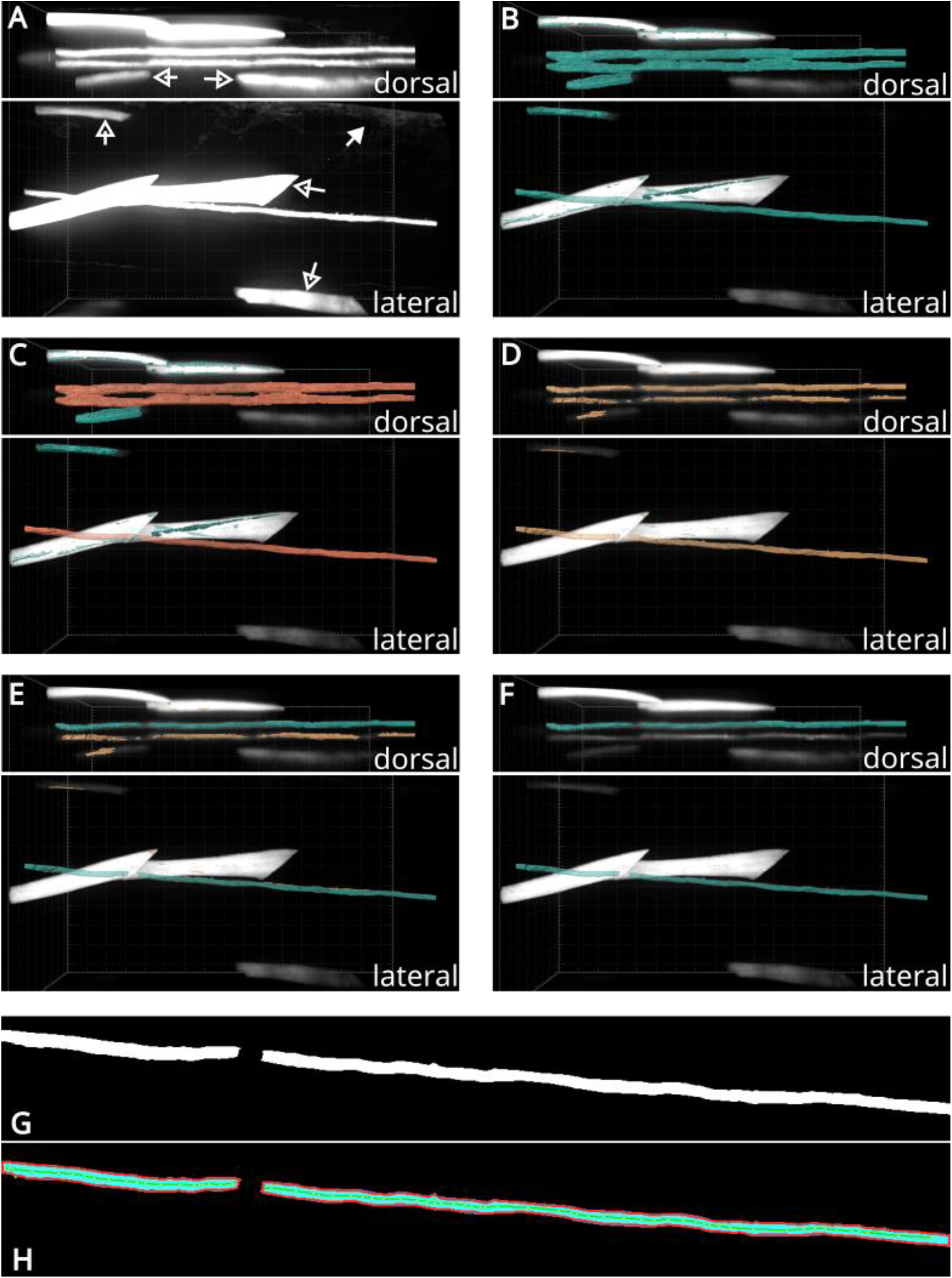
Creating an automated image analysis pipeline to measure axon diameter changes: Stepwise object detection and processing. Front and top 3D views of z-stacks rendered in Arivis Vision4D showing: A) Overly contrasted greyscale look-up-table to highlight problematic skin reflection/autofluorescence (solid arrow), and ectopic muscle expression of GFP (hollow arrows). B) Low-resolution ML model trained to exclude skin and muscle leaving candidate axonal objects (green). C) Object filtering to exclude small objects (green), leaving target Mauthner axons (orange). D) High-resolution ML model trained to tightly segment clear and reliably measure candidate axons (yellow). E) Object size filtering followed by a clustering python script determines objects most likely to be the Mauthner axon closest to the objective (green), other objects (yellow) are discarded. F) Final segmented Mauthner axon (green) used for measurement. G) Mauthner axon object slices are exported and z-projected to create a 2D planar mask. H) 2D planar mask is polygonised and the polygonal skeleton used to take multiple skeletal width measurements across the length of the detected axon. A-H) minor grid ticks are 5 µm, major grid ticks are 20 µm.

Many image analysis platforms, both open-source and commercial, now implement machine learning (ML) or Artificial Intelligence (AI)-based training and segmentation. There are many image analysis platforms both commercial and freeware. Freeware platforms offer flexibility and customization while commercial platforms often focus on reduction of barriers to high-quality image analysis. Both are valid and useful tools. Here, we employed multiple levels of random forest pixel classification using Arivis Vision4D to generate an automated and objective pipeline for consistent batch measurement of Mauthner axon diameter in live zebrafish. Commercial platforms like Vision4D, Imaris, and others provide a GUI and user-friendly method of training models and implementing pipelines of processing algorithms. The software used here (Vision4D) allows users to see the results of their changes and build pipelines piece by piece in a single environment. Models can be exported and reused, and when mismatched parameters or incorrect application of a model is attempted, it is flagged with warnings and pointers on how to correct. Similar functionality now exists in GNU General Public License software Ilastik and the models from Ilastik can be imported into Vision4D and other image handling packages like ImageJ.

#### Model Training

Single field imaging of the spinal cord at somite 15-16 was conducted as described above (see also materials and methods). Selected confocal stacks from multiple runs, encompassing multiple batches, were imported into Vision4D to create a training data set (alternative software: ImageJ/Ilastik). Each slice of each image was heavily blurred (gaussian, 146µm radius, alternative software ImageJ) to create an additional low-resolution denoised channel. This channel and the original were used to train a low-resolution random forest pixel classifier that would detect, very roughly, the Mauthner axons and exclude most artefacts or unwanted signal (highlighted in Figure 2A-C, muscles (hollow arrows), autofluorescence through skin (solid arrows)). Next, a high-resolution model was trained using only the original channel raw data (Figure 2D). To achieve sharp segmentation of axons, without aberrant object growth and inclusion of neighboring artefact objects, it was necessary to add more negative classes. Signal loss at depth from scattering and absorption, as well as biological issues like dimming of axonal signal at somite edges, and striation patterns present on Mauthner axons and muscles, meant that it was necessary to train the ML model to detect the clearest axon segments (i.e. those closest to the objective lens) and those free from occlusion by fluorescent muscle or other artefacts. This results in poor segmentation of the axon furthest from the objective lens, and often broken detection of the axon where it underlies the edges of somites (Fig 2D).

Both models are imperfect and lead to fragments of artefacts being classified as candidate ‘axon’ objects: a result of the limitation of random forest pixel classifier models and of minimising the amount of training data. However, combined with logical flow of object filtering and object math, incorrectly classified objects were eliminated (Fig 2C). To segment untrained images, the models and logic flow were combined into pipelines for batch processing (Figure 3A). The batch process was created using incrementally tested values, including object filters, object math, and object reshaping, into a single GUI pipeline (steps 1-9 in workflow diagram Figure 3A). Because the zebrafish larvae were positioned laterally the Mauthner axons lay at different z-depths within the tissue. Therefore, the imaging and segmentation quality could vary depending on the positioning of the fish and proximity to the objective lens. To account for this, we divided the axons into “top” and “bottom” groups based on their relative z-position, enabling separate analysis of the better-resolved (top) axon. To sort the axon objects into those nearest the objective lens (top) and those furthest (bottom), relative or relational object filtering was required (e.g. “highest” of two objects with “similar” x centroid values, or group objects with “similar” z centroid). Vision4D lacks any relative object comparators natively and does not support calculated object variables. It does support Python scripting via an application programming interface (API) and so a python script was created to cluster objects in the z-dimension and sort the clustered objects into “top” and “bottom” axons (imaged lateral-side-up) (Figure 2E & F). This step works partially outside of the image handling environment and so is represented as an additional process (Figure 3A, dotted arrows). If the project was handled in ImageJ, a java script could be used at this point to achieve the same goal.

**Figure 3:**
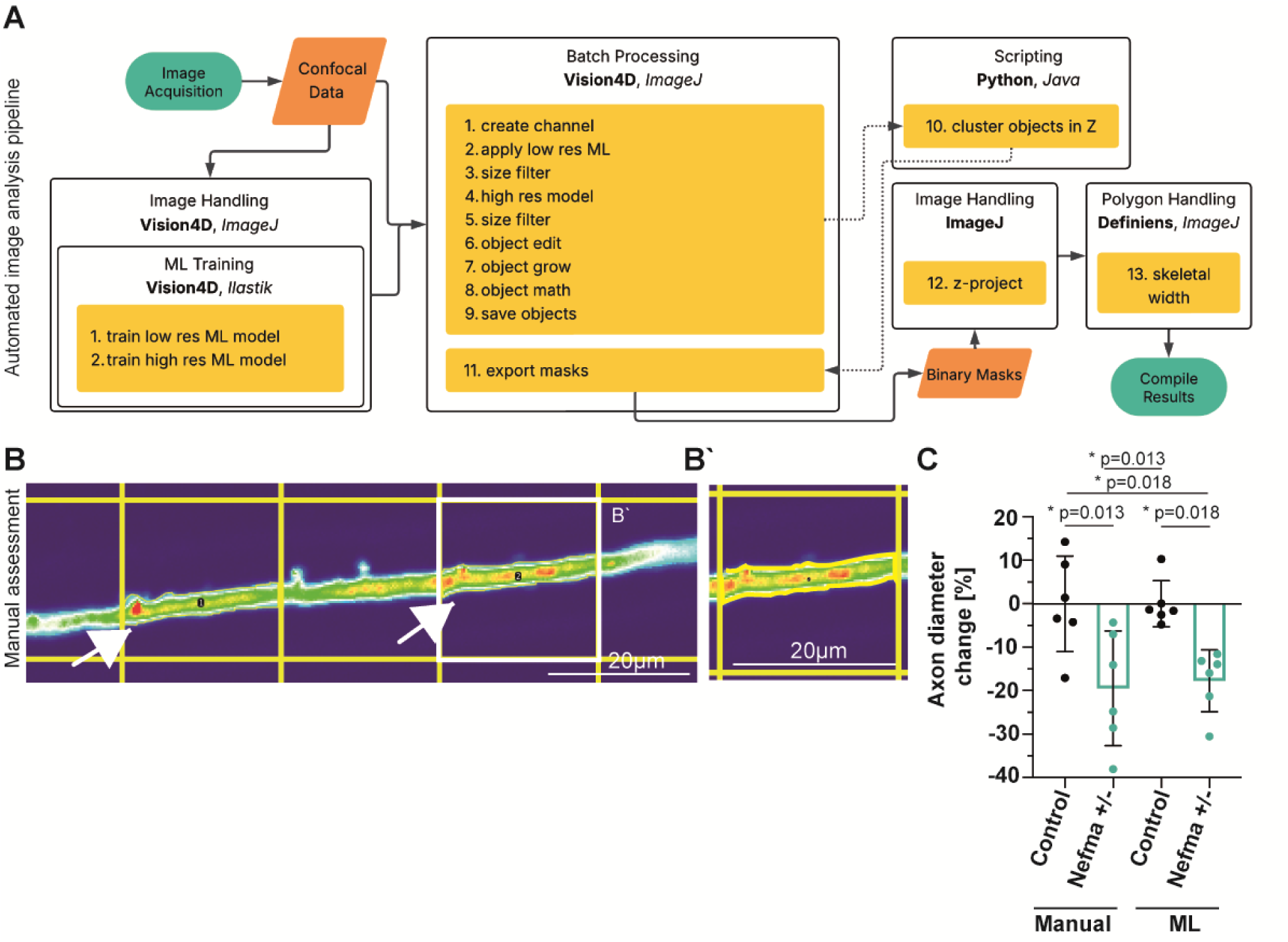
Manual assessment confirms reliability of the machine learning-based quantification of Mauthner axon diameter. A) Workflow diagram illustrates the step-by-step process to create our fully automated image analysis pipeline, the tools used in our study are in bold and alternative suggestions and open-source tools are indicated in italic. B) Representative thermal look-up table (LUT)-thresholded image of a Mauthner axon with an overlaid random measurement grid (500 µm² spacing). Two selected grid squares intersecting the axon are highlighted to indicate regions where axon diameter was manually traced (arrows). B′) Zoomed-in view of a square 2 showing manual tracing of the Mauthner axon outline (turquoise pixels) using polygon selection. Manual measurements were performed on at least five regions per sample to calculate mean axon diameter, as described in Methods. C) Comparing the normalized percentage change in Mauthner axon diameter between control and a mutant with smaller Mauthner diameter (Nefma+/-) shows that both manual assessment and automated analysis detect a similar, statistically significant decrease. One-way ANOVA with multiple comparison test was used to assess statistical significance. Error bars represent mean with SD. P-values are presented as * p<0.05. Scale bar=20µm.

A limitation of our chosen image handling platform was its lack of comprehensive object measurements. For our purposes, we needed to measure the width of the top axon across its length. An applicable method is to measure the polygonal skeletal width across the length of the object. To obtain an average width measurement for each axon, slice masks for the top Mauthner axon were exported, projected in Z, and measured in a commercial package known to have this functionality (Definiens DeveloperXD). An example of the process can be seen in Figure 2G-H. The workflow diagram (Figure 3A) shows the steps required to create our fully automated image analysis pipeline, states the tools used in this study (bold text), and suggests alternative or open-source tools that can achieve the same goal (italic text).

To test the performance of the automated image analysis pipeline, we used an available transgenic line^50^ that we found to exhibit smaller Mauthner axon diameters and imaged it using our standardized VAST-SDCM settings. We then compared axon diameter measurements obtained from the automated analysis pipeline with those from fully manual assessments (see Material & Methods and Figure 3B and B‘). Both methods show a significant decrease in Mauthner axon diameter in the reporter line compared to controls (Figure 3C). Taken together, we reasoned that our imaging and analysis pipeline is sensitive enough to detect changes in Mauthner axon diameter.

### Chemical screen reveals compounds that change Mauthner axon diameter

After establishing our imaging and analysis pipeline, we next wanted to carry out the screen to identify factors that changed axon diameter. We used the LOPAC^1280^ library because its well-described compounds, with annotated targets that represent diverse pathways, with a particularly high portion related to neurotransmission- and ion channel-function. Because it was previously shown that axon diameter can change upon neuronal activity ex vivo^9^, we reasoned that this library might identify candidates to testing the hypothesis that activity impacts axon diameter growth in vivo.

For zebrafish screens, compounds in the LOPAC library have commonly been used at around 10µM to prevent severe impacts on health^51,52^. Since we treat our larvae at slightly later stages (3-5dpf) we initially tested 30 randomly chosen drugs at 2, 10, 50, and 100µM and assessed the survival and health of fish. At 10µM we observed little impact on health and survival of larvae (∼1%) and thus selected 10µM as the primary concentration for each compound. Compounds that did cause larval death at 10µM were rescreened at 2µM in the primary screen. To ensure that our screen would detect subtle changes of 10%, such as those identified using rapamycin, we decided to use n=9 animals per compound. To maximize our throughput and test more compounds per 96-well plate, we placed 3 fish/well which allowed testing of 30 drugs per plate (9 animals per drug): each 96-well plate of 3 animals per well was imaged in ∼5 hours. To accommodate for clutch specific differences in axon diameter and for any changes that may occur to diameter over the imaging period, we included 3 wells with 3 fish/well treated with 0.5% DMSO as clutch specific controls, spread out across the 96-well plate. Importantly, DMSO at 0.5% does not impact axon diameter growth. In the screen animals were treated with compounds from 3-5 dpf, imaged at 5 dpf and assessed using our automated analysis pipeline (Figure 4A). The resulting Mauthner diameters for each image were matched to their respective treatment using custom-made python scripts that matched the time and well-based information of each image (clutch, drug treatment) with the measured diameter values. To assess how test compounds influence axon diameter, we normalized compound effects on axon diameter growth relative to controls and calculated a z-score to rank how compounds differed from controls. We did so by comparing axon diameter measurements for each drug-treated sample to those of the corresponding DMSO-treated clutch and plate-specific control (see Methods for calculations). This standardization is common practice in high-throughput drug screening approaches because it accounts for variability across wells, plates and batches and allows ranking of the impact the identified hits have. Given that we image at subcellular resolution and aim to detect changes as small as 10% in an axon as big as the Mauthner axon, we set an artificial threshold of z-score > +1.2 or < –1.2 to define hit compounds in the primary screen. To minimize technical, imaging, or analysis-related errors we manually inspected the z-stacks and masks for each candidate hit as well as their respective controls. In some cases, imaging errors (e.g. drift of fish in z) led to incorrect detection of the Mauthner axon. Therefore, we excluded any data points without accurate axon identification and mask-based measurement from our analysis (For a detailed description of each step, see Materials & Methods).

**Figure 4:**
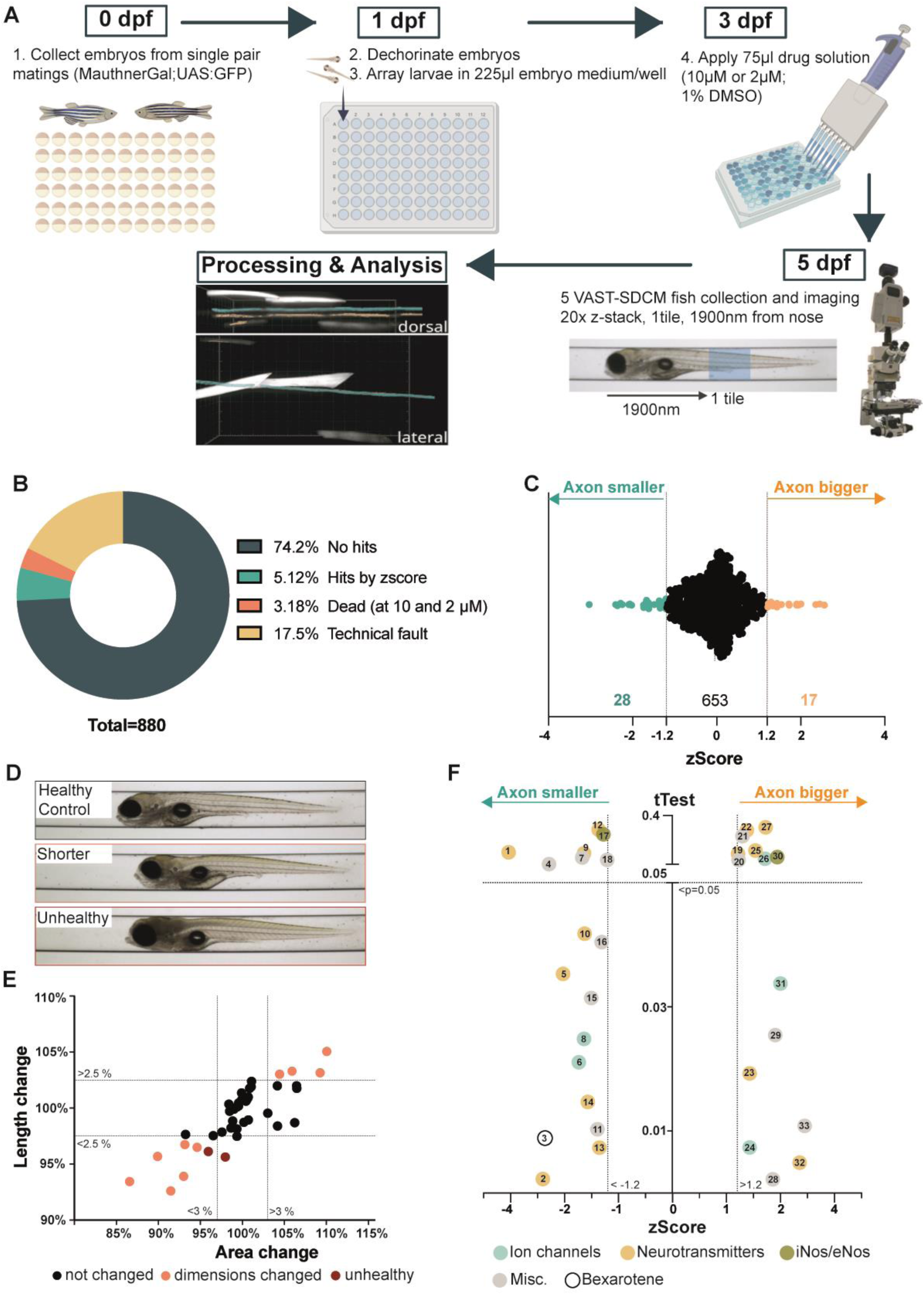
Phenotypic screen identifies compounds that change Mauthner axon diameter. A) Schematic illustrating the workflow of the screening pipeline. B) In the primary screen 880 drug compounds were tested at either 2µM or 10µM for their potential to alter Mauthner axon diameter and approximately 5% were initially identified as hit compounds. A further 3% resulted in death at both dilutions and 17.5% were excluded due to technical issues. C) The z-Score was calculated by comparing each drug with its respective DMSO-treated clutch-specific control. A z-Score of >1.2 or <-1.2 was considered changed in diameter. 28 compounds decrease and 17 compounds increase Mauthner axon diameter. D) Brightfield images show a healthy appearing control (top), a larva with smaller body dimensions (middle) and one with an unhealthy appearance (bottom). E) Body dimensions (length and area) of each hit compound and their respective DMSO-treated clutch control were subsequently assessed from brightfield images. Changes in over 3% body area and 2.5% length were considered as exclusion criteria (pink). Toxic compounds are indicated in red. F) Results of the primary screen displays hit compounds ranked by z-Score (X-axis) and statistical assessment using Students t-Test (Y-Axis) comparing each hit compound with its DMSO-treated clutch specific control. 33 candidate hit compounds are color coded by class of action (turquoise Ion channels, yellow Neurotransmitters; green iNos/eNos species and grey Miscellaneous). In total, 18 drug compounds decrease Mauthner axon diameter beyond z-Score threshold of −1.2 and 15 drug compounds increase Mauthner axon diameter above threshold of 1.2. See also Table 1 for description of compounds.

In total, we tested 880 drugs for their ability to affect axon diameter. We found that 28/880 drugs (3.18%) were lethal at both 10 and 2 µM dilution and 154/880 treatments (17.5%) had to be excluded due to technical difficulties (low image quality, false detection of the Mauthner, n number<4). Of the remaining compounds, 653 (74.2%) did not meet the z-score threshold, while 45/880 compounds – approximately 5% – were classified as hit compounds (Figure 4B and C).

To check for the impact each candidate hit might have on the health of the fish, we manually assessed all accompanying RGB photographs acquired by VAST, for any discoloration (sign of ill health) or deformations. We excluded 2 compounds because fish were discolored or their spine slightly deformed (Figure 4D). Next, we wanted to make sure that the remaining 43 hit compounds did not influence axon diameter secondary to impacting the overall size of the fish. To exclude candidate hits that primarily impact body dimensions we measured body area and length of all animals treated with candidate hits and their respective controls. Determining relative subtle changes to animal size (e.g. on the order of 10%, Figure 5A) by eye is very challenging so we applied further automated methods. This time, we trained a random forest pixel classifier to recognize whole fish larvae (Figure 5B). We measured body area and length using measurements built into DeveloperXD as described in materials and methods (Object Area, Length (bounding box)). To define exclusion criteria, we first assessed variation in whole-body area and fish length. We did so by pooling the values for area and length of DMSO-treated controls across various clutches and imaging rounds and calculating their mean values, and standard deviation. A single standard deviation from the mean length of all controls was 2.25% and of area was 2.92%. Therefore, we considered changes exhibited by hits that were greater than a single standard deviation as likely affecting animal size and thus set a change of both 2.5% in length and 3% area as an exclusion threshold for altered body dimensions (see Material & Methods; stippled lines Figure 4E). This resulted in the exclusion of 10 hit compounds (Figure 4E). Validating these exclusion criteria, we found that changes in body area and length were significantly correlated with axon diameter for the 10 excluded hits (R^2^=0.72 for area and R^2^=0.69 for length) (Figure 5C). In contrast, there was effectively no correlation between to axon diameter and body area or length for the remaining 33 candidate hits (R^2^=0.02 for area and R^2^=0.01 for length) (Figure 5D), suggesting specific alteration to axon diameter independent of body size.

**Figure 5:**
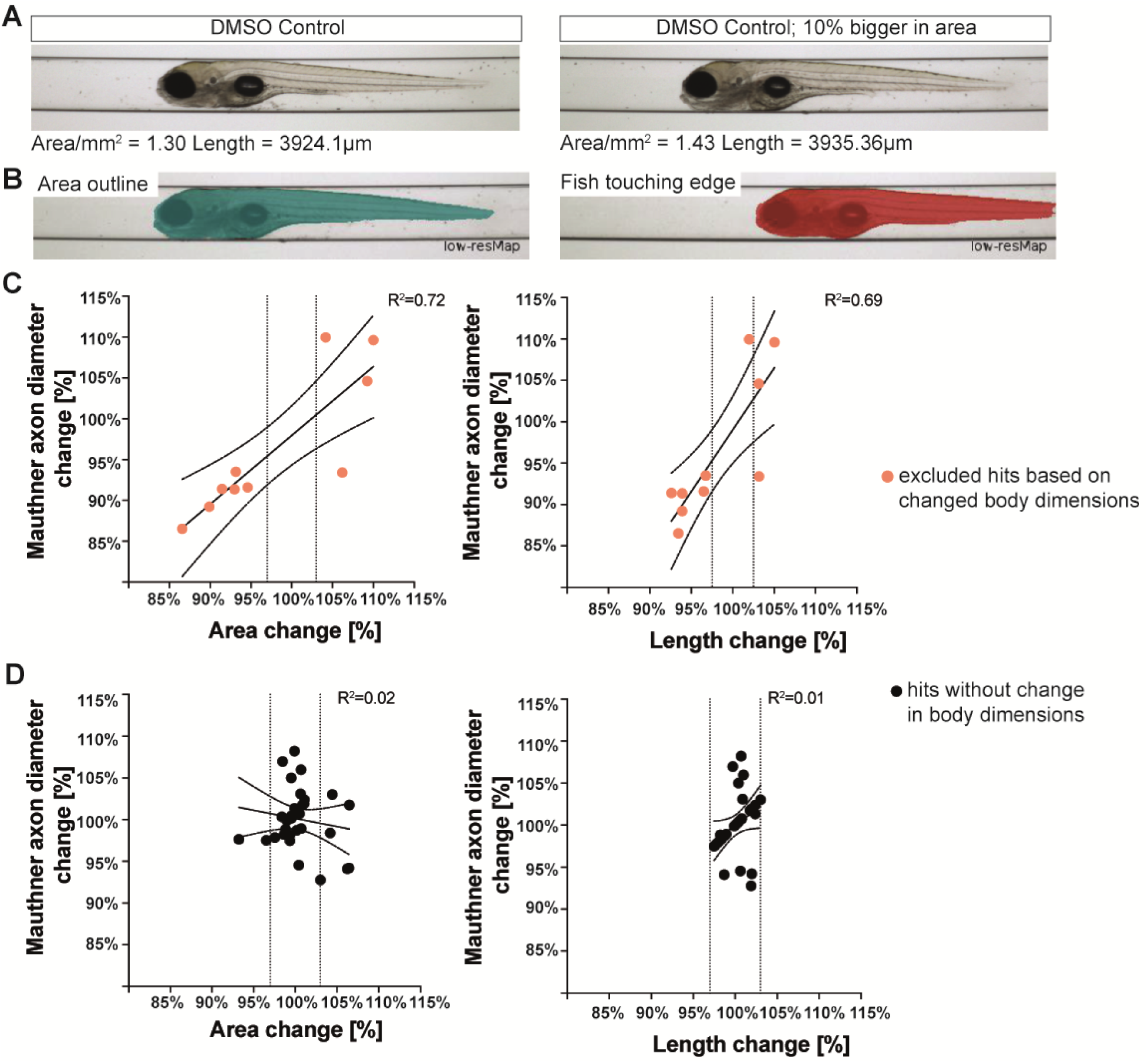
Assessment of exclusion criteria for body dimensions. A) The two depicted brightfield images acquired on VAST-SDCM of DMSO treated controls vary by 10% in area which we cannot manually assess by eye. B) Body dimension (length and area) was calculated on brightfield images from VAST-SDCM by training a machine learning classifier in DeveloperXD. Exclusion criteria (3% area change and 2.5% length change) were estimated based on averaged variations across different DMSO-treated controls (See Methods for details). C) Hit compounds that were excluded based on body size are shown in pink. For those, both area and length correlate with axon diameter (R^2^=0.72 area and R^2^=0.69 length). D) Hit compounds without change in body dimensions are shown in black (right) and have no correlation between area or length and axon diameter (R^2^=0.02 area and R^2^=0.01 length).

Among the 33 remaining hit compounds, 18 decreased Mauthner axon diameter, while 15 increased it (Figure 4F, Table 2). Notably, 19 of the overall 33 hit compounds are linked to neurotransmission and ion channel function. This enrichment is consistent with the LOPAC library composition, in which ∼60% are neurotransmission and ion channel related compounds. In contrast, other compounds target nitric oxide species, lipid signaling, phosphorylation, cytoskeleton, and extracellular matrix. To complement the z-score ranking, we performed a Student’s t-test comparing each hit compound to its respective controls, enabling both standardized effect size assessment and statistical validation. This analysis identified 18 out of 33 compounds that significantly alter axon diameter. Table 2 highlights all hit compounds, including their ranking metrics and action class.

**Table 2.**
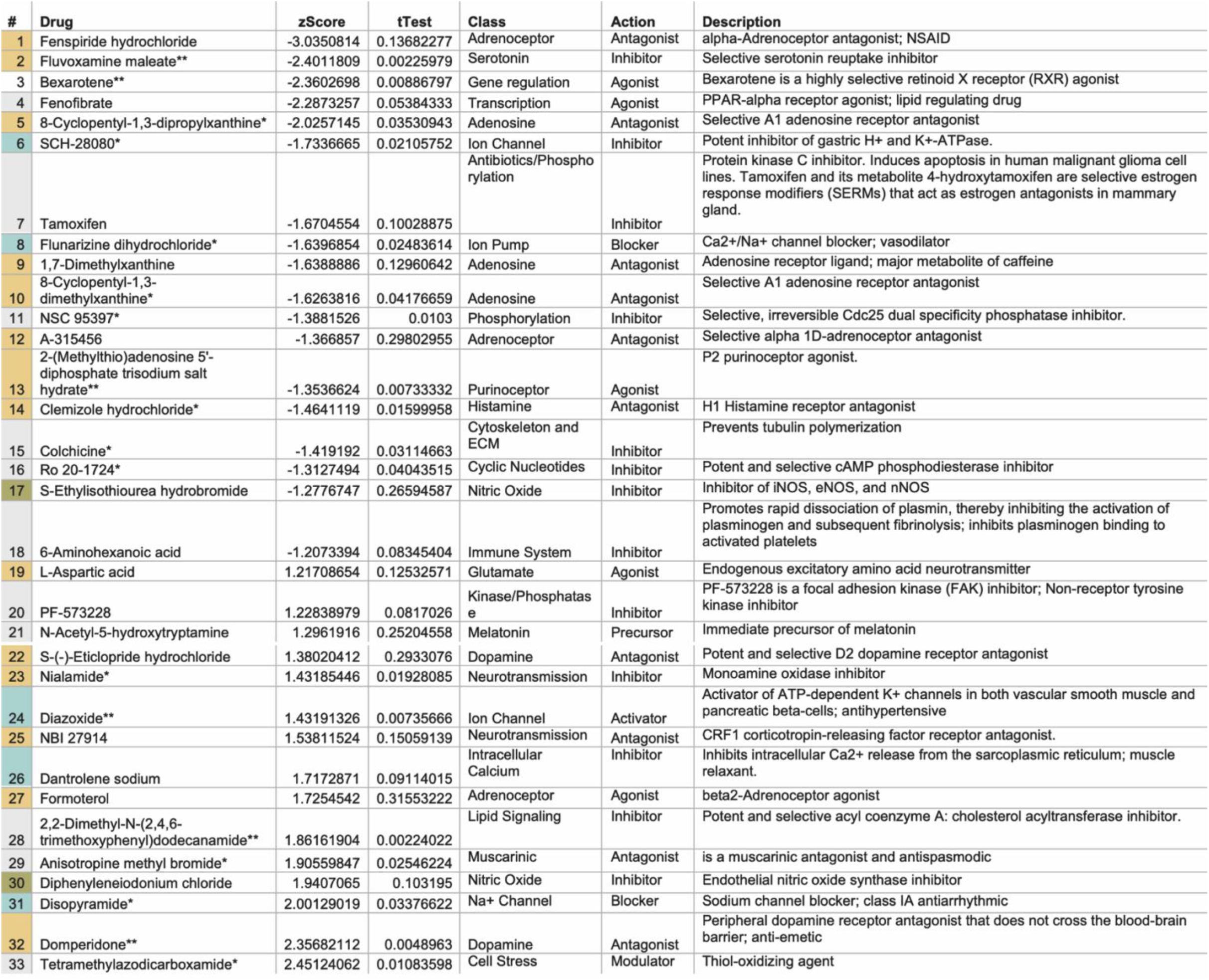
List of hit compounds that change axon diameter. Complete list of candidate hit compounds, putative targets and description, that change Mauthner axon diameter ranked by zScore. The z-score was calculated comparing the diameter for each compound with values of respective DMSO-treated clutch specific controls. Student’s t-Test was used to additionally assess statistical significance between both groups. Compounds that change overall body dimensions or overall health are excluded. Class, action and description are shown for each identified hit compound.

In summary, we performed the first, to our knowledge, in vivo discovery screen designed to identify compounds that modulate axon diameter. Out of 880 compounds tested, following rigorous data curation, we identified 33 compounds that influence Mauthner axon diameter growth. We provide our complete list of hit compounds to the neuroscience community as a resource to facilitate further investigations.

### Validation of hit compounds reveals that beta-2-adrenoceptor agonists and dopamine antagonists increase Mauthner axon diameter growth

Next, we wanted to validate hit compounds from our primary screen. We first assessed the action class of compounds and prioritized hit compounds that might influence axon diameter growth by modulating neuronal activity. In addition, we prioritized compounds according to 1) their z-score and effect size rank; 2) repeated appearance of their class of action amongst hits; 3) presence of both antagonist and agonist of the same class that introduce opposing effects on diameter; and 4) their ability to increase the diameter of the already large Mauthner axon. Based on these triage steps we first focussed on compounds linked to adrenoceptors (2 antagonists decreasing and 1 agonist increasing diameter) and dopamine signaling (2 antagonists increasing axon diameter).

We ordered the same compounds or those that target the same pathway from different suppliers. To assess dose-dependent effects of the new compounds on Mauthner axon diameter we performed serial dilutions ranging from 1-100µM and manually assessed compound toxicity by eye to determine at which concentrations larvae induced developmental deformities or lethality. We excluded these concentrations from further analysis. When available, we re-tested the original LOPAC library compound at the concentration we used in our initial screen (10µM or 2µM). While the VAST-SDCM system and our imaging pipeline enabled us to perform the large-scale primary screen, our validation experiments did not require high-throughput imaging. To achieve higher resolution imaging, we reverted to our previously established method for imaging axon diameter by confocal super-resolution imaging (Figure 6F and F‘). We imaged Mauthner axons of drug-treated and DMSO-treated control larvae and used the semi-automated script to measure axon diameter, given that our ML-driven image analysis was bespoke to images acquired on our VAST-SDCM platform. To account for clutch-specific variability we normalized axon diameter by calculating the change relative to DMSO-treated controls of the same clutch imaged within the same time window of ∼3h.

**Figure 6:**
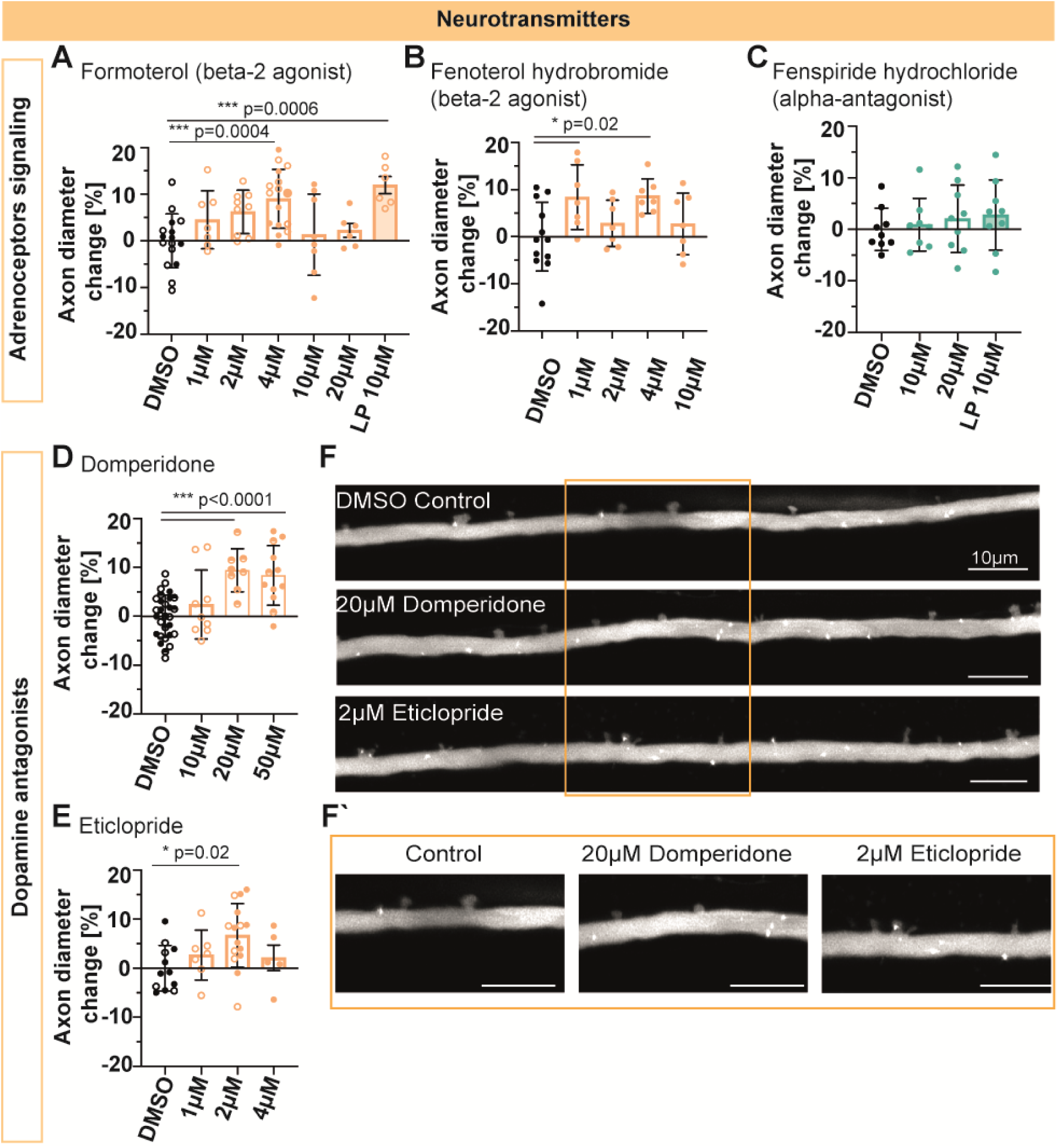
Beta-2-adrenoceptor agonists and dopamine antagonists increase Mauthner axon diameter growth. Validation of selected hit compounds different vendors was performed at different concentrations (1-50µM) and compared to control fish treated with 0.5–1% DMSO. Fatal or health impacting dilutions are not shown. The change of Mauthner axon diameter for each drug was normalized to the clutch specific DMSO-treated control and calculated in %. If available, the original LOPAC1280 drug was tested at its original dilution (LP). A) Beta-2 Adrenoceptor agonist Formoterol resulted in a significant increase in Mauthner axon diameter at 4µM and when testing the original LP dilution at 10µM. B) A second beta-2 adrenoceptor agonist, Fenoterol hydrobromide shows an increase in Mauthner diameter at 1µM and 4µM. C) Alpha Adrenoceptor antagonist Fenspiride hydrochloride did not show the initially detected decrease in Mauthner axon diameter, no significant change was detected. D and E) Both Dopamine antagonist Domperidone and Eticlopride resulted in a significant increase in Mauthner axon diameter. F – F‘) Fish were imaged using 40x objective at a Zeiss LSM880 with AiryScan FAST mode with 1.8 zoom for validation studies. Confocal images and zoom in (F‘) visualize increase in Mauthner axon diameter when fish were treated with Dopamine antagonists Domperidone and Eticlopride. Individual clutches are shown in open versus closed circles. One-way ANOVA followed by Dunnett‘s multiple comparison test was used to assess statistical significance. Multiple adjusted p-values are shown in each figure panel and are presented as * p<0.05, ** p<0.01, *** p<0.001. Error bars represent mean with SD. Scale bars=10µm.

Our validation tests revealed that the beta2-adrenoceptor agonist Formoterol increases the axon diameter of the Mauthner at 4µM, with the original LOPAC1280 compound also showing a significant increase in diameter at 10µM (Figure 6A and C). To assess another beta-2 adrenoceptor agonist, we applied Fenoterol hydrobromide and detected a significant increase in axon diameter at 2µM and 10µM (Figure 6B and C). We did not detect changes in Mauthner axon diameter when applying the alpha-adrenoceptor antagonist Fenspiride hydrochloride, which decreased axon diameter in the initial screen (Figure 6D). We were not able to acquire the third adrenoceptor-related compound A315456 on the market. Next, we assessed the two hit compounds targeting Dopamine signaling: the dopamine antagonists Domperidone and Eticlopride. Both compounds significantly increased diameter in our validation studies (Figure 6D-F‘). Taken together, we have validated two hit classes, beta-2 adrenoceptor agonists and dopamine antagonists, as affecting axon diameter growth.

## Discussion

Zebrafish are a powerful model for in vivo discovery screens, with ongoing innovation in biosensors and reporters, imaging and image analysis, now making it possible to investigate the biology of previously intractable processes ^47,51–55^. Here, we have brought the power of scalable high-resolution high-content imaging of zebrafish to bear on the biology of axon diameter, a largely overlooked, but essential feature of neuronal morphology and neural circuit structure and function.

To screen for potentially sub-micron scale changes in axon diameter we needed 1) sufficient imaging resolution to detect changes to axon diameter, 2), scalability required to image and analyze a sufficiently large number of compounds to power discovery potential; 3) sufficiently robust analyses to limited false-positives and false-negatives findings. Therefore, we used the Mauthner neuron ^44–46^ as a model, because its axon grows to ∼5µm in diameter by 5dpf^44^ meaning that changes could be resolved using our automated confocal screening system (VAST-SDCM). We tested 880 compounds for their potential to alter axon diameter and through a robust automated analysis pipeline identified 33 hit compounds that changed Mauthner axon diameter. Our validation studies confirmed the effects of some hit compounds (discussed below), but also revealed false positive hits, a common occurrence in discovery screens. In the context of our screen a number of points are noteworthy: 1) automated segmentation of axons from acquired images was not 100% accurate, meaning that a large number of samples were discarded such that the n number for certain compounds will have been lower than hoped for, contributing to both false positive, and likely also false negative hits and 2) chemical screens can be subject to variability due to deviation from expected compound concentrations, plate effects such as evaporation and edge effects, meaning that rigorous validation steps, as we have outlined are required. Nonetheless, our screen and validation pipeline has already confirmed hit compounds that do affect axon diameter, namely those affecting dopamine signaling and adrenoreceptors.

Domperidone and Eticlopride are both D2-dopamine receptor antagonists. Domperidone has been shown to affect locomotor behaviour in zebrafish with D2-receptor antagonism associated with increased neuronal excitability across vertebrates ^56–58^. Interestingly, dopaminergic neurons have been shown to modulate the sound-evoked Mauthner cell mediated C-startle response^59^. This suggests the possibility that systemic dopaminergic manipulation may influence the activity of the escape circuit, including Mauthner neuron activity, which may in turn influence axon diameter.

Beta-2 adrenoceptors are G protein–coupled receptors that mediate various physiological responses, including smooth muscle relaxation and subsequent vasodilation^60,61^. Since beta-2 adrenoceptors can be activated by endogenous catecholamines like noradrenaline and adrenaline, which are known to be linked to changes in locomotion, they may also exert an effect on axon diameter via modulating neuronal activity. We know from ex-vivo preparation of hippocampal neurons that axons can widen upon high-frequency stimulation^9^. However, whether axon diameter changes upon neuronal activity has not been shown in vivo. If the changes in axon diameter induced through our hit compounds reflect alteration of neuronal activity and circuit function or affect alternative mechanisms that influence axon diameter remains to be seen. In addition to deconstructing how our currently-validated hits affect axon diameter, extensive future studies will be required to test and confirm the activity of the other hits from our screen. This will allow to expand our understanding of the many mechanisms likely to influence axon diameter, including those suggested by our screen, such as lipid signaling, pathway-specific phosphorylation, and ECM/cytoskeletal regulation.

Our study represents the first dedicated effort to discover the molecular basis of axon diameter growth but was limited to the investigation of one neuronal subtype at one developmental age. Therefore, it will be very important that future follow-up studies characterize the effect of hit compounds on different neurons that have axons of distinct sizes, and at distinct stages of system and circuit maturation. In addition, to assess the molecular mechanisms underlying compound induced changes will require complementary genetic perturbations of candidate pathways. Furthermore, it will be important to extend discoveries made in zebrafish to mammalian, including human models, both in vitro and in vivo. In summary, we hope that our screen serves as foundation to help further understanding of the biology of axon diameter in vivo.

## Methods

### Zebrafish husbandry and transgenic lines

Adult zebrafish were housed and maintained in accordance with standard procedures in the Queen’s Medical Research Institute zebrafish facility at the University of Edinburgh. All experiments were performed in compliance with the UK Home Office, according to its regulations under project licenses PP5258250 and PP3290955. Adult zebrafish were subject to a 14/10 hr, light/dark cycle. Embryos were produced by pairwise matings, collected within 2 hours post-fertilisation (hpf) and raised with 50 embryos per dish at 28.5°C in 10 mM HEPES-buffered E3 Embryo medium or conditioned aquarium water with methylene blue. All experiments used zebrafish larvae up 5 days post-fertilisation (dpf) on a wild type (AB/WIK/TL) or nacre^−/− 62^ background. At these ages, sexual differentiation of zebrafish has not yet occurred. The following transgenic lines were used for this study: Tg(hspGFF62A:Gal4)^48,63^, Tg(UAS:GFP)^48^, Tg(nefma-mcherry)^50^. When characterizing the Tg(nefma-mcherry) transgenic reporter, we noted a reduction in Mauthner axon diameter in the heterozygous form (data not shown) and used it as a reporter for smaller axon diameters. Controls refer to clutch-specific siblings.

### Hardware setup (VAST-SDCM platform)

All animals in the primary LOPAC screen were imaged using the VAST-SDCM platform described in Early et al^47^. Briefly, three fish were loaded into each well of a 96 well plate and brought to a CSU-X1 spinning disk (Yokogawa) unit with dual AxioCam (Zeiss) 506 mounted on an Examiner frame (Zeiss) using a combination of LP Sampler and VAST robotic fluidics (Union Biometrica). We used a W Plan-Apochromat 10x/0.5NA M27 75mm dipping lens (Zeiss) and generated stitched z-stacks for whole fish overview images (Figure 1A). Otherwise, for axon diameter measurements, the system was configured to acquire a single z-stack 1900µm distal from the tip of the larvae head using a W Plan-Apochromat 20x/1.0NA Corr DIC M27 75mm objective lens (Zeiss). The single 333 x 249 x 300 µm (XYZ) stack with 2×2 binning (286nm pixels) and 300ms exposure was used for optimal speed while maintaining signal:noise ratio for image processing.

### Chemical screen: larvae preparation and compound treatments

Out-crosses of the Tg(hspGFF62A:Gal4,UAS:GFP) transgenic line to nacre fish were used for the primary screen. Embryos were collected within 2 hpf and raised with 50 embryos per dish. Clutches were separated. Embryos were enzymatically dechorionated at 24-30 hpf using protease from Streptomyces griseus (0.5 mg/mL for 6 min) (Sigma-Aldrich, St. Louis, MO) and washed with E3 media. Afterwards, three embryos per well were manually arrayed into 96-well plates in 225 μL E3 media. At 2dpf and 3dpf, larvae were checked for viability; unhealthy animals were manually removed and replaced in equal volumes with clutch-matched larvae. First, 10 mM compound stocks in DMSO (LOPAC^®1280^, Plate 1-11; Sigma-Aldrich, St.Louis, MO) were serially diluted using a multi-channel pipette to 2mM, 100% DMSO and frozen at −80°C. On the day of drug treatments, a 4X concentrated treatment solution was generated by diluting 5µl or 1µl in 245 µl E3 media (40µM and 8µM respectively, 2% −0.4% DMSO). Between 70-75 hpf, 75 µL of this 4X stock was added directly into the larval wells for a final concentration of 10 µM or 2 μM in 0.5-0.1% DMSO). Each compound was tested on 3 wells of 3 fish per well (n = 9) and compared against 0.5% DMSO-treated negative controls of the same clutch (n=9-12) within the same plate. Treatment plates were incubated under standard temperature conditions for 2 days without compound refreshment. At 5 dpf, larvae were anaesthetised with 600 µM tricaine before imaging. One 96-well plate with 30 compounds and respective controls was imaged within 4.5-5.5 h. The control wells were spread out over the plate and/or imaged throughout the day to spread control variability due to development whilst imaging live animals over time.

### Automated image analysis pipeline

A machine learning training set that covered the range of sample and image variation encountered in the study was created in Arivis Vision4D v3 from full XYZ stacks in CZI format acquired from 9 independent imaging runs. Two random forest pixel classifiers were created using the “Machine Learning Trainer” feature: LowResML and HighResML. Both were set to use 100% scaling as well as all features at all pixel sizes. Training samples were annotated on 37 images in this data set as follows: LowResML - Axon class (71 samples, positive class), Fuzzy class (271 samples, negative class), Background class (139 samples, negative class); HighResML – AxonHighRes class (88 samples, positive class), fuzzyHghRes class (461 samples, negative class), muscleHighRes class (72 samples, negative class), background class (166 samples, negative class). Both models were required sequentially to successfully detect target Mauthner axons at full resolution. LowResML was used to detect target Mauthner regions and exclude most of the skin autofluorescence and ectopic muscle GFP expression (Figure 2A & B). Following size and shape filtering of low-resolution segmented objects (Figure 2C), highResML was applied (Figure 2D). High resolution segmented objects were only kept if they were of the correct AxonHighRes class and were contained within the low-resolution Axon class (Figure 2E). High resolution axon segments were then sorted, using a Python script linked to Vision4D, into upper, lower, and solo Mauthner axons (Figure 2F). Single-plane binary masks were exported and z-projected in FIJI before being imported into Definiens DeveloperXD for measurement of polygonal skeletal width.

The detailed workflow is comprised of the following steps and depicted in Figure 3A:

1. Create additional gaussian channel

1. Apply low resolution ML model
2. Filter objects for size (>16000µm^2^)
3. Apply high resolution ML model
4. Filter objects for size (>13000µm^2^)
5. Fill inclusions (fill holes) in low-res objects
6. Dilate low resolution objects (5 pixels)
7. Object math: intersect low resolution objects with high resolution objects
8. Keep only intersected objects
9. Sort objects into Z-dimension clusters and keep top-Z clusters (integrated python script)
10. Export masks of remaining Mauthner objects slice-wise
11. Z-project binary masks to 2D image
12. Import, segment, and measure binary masks

After individual values for Mauthner axon diameters were measured a python script was used to filter and sort axon diameter values according to image day, drug treatment and clutch. Afterwards, the Z-Score was calculated to rank drugs and detect each drug’s effect on Mauthner diameters of drug-treated fish. The following equation was used

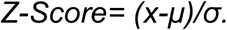

Here, x is the mean of the measured axon diameters from each drug; µ is the mean of the axon diameter from the clutch and plate specific, DMSO-treated control and σ is the standard deviation of the respective control group. A z-score > +1.2 or < –1.2 was defined as hit compound in the primary screen. Afterwards, Mauthner axons detected in Vision 4D and axonal mask from Definiens for each hit compounds and respective control were manually assessed for 1) optimal imaging (for example no drift in z), 2) correct detection of the Top Mauthner and 3) correct mask generation. If any of them did not guarantee optimal detection and tracing of the Top Mauthner axon the values were excluded. If for a given compound the n-number is lower than 4 we excluded that compound as well. All excluded compounds fall under the category technical difficulties (Figure 4B).

### Whole zebrafish size dimensional analyses

Colour photographs from VAST-SDCM were used to assess the health and overall dimensions of fish treated with hit compounds according to z-Score. Images of each fish were imported into Definiens DeveloperXD image analysis software (Definiens AG). A montage was created from 20 images and used to train a random forest pixel classifier to detect fish photographed inside the capillary tube. The trained machine learning classifier was used in an automated DeveloperXD pipeline to detect all fish, exclude those touching the edges of the photograph, and then measure the Area & Length (bounding box) of each fish using built-in functions with those names. The overall appearance of the fish was assessed manually.

To estimate variations in length and area, both values of DMSO-treated control fish across 14 different clutches and 10 individual VAST-SDCM runs were averaged. We considered this spread in controls as the normal change in dimensions. For exclusion criteria the single standard deviation from the mean area change (2.92) and length change (2.25) were calculated. Changes greater than the single deviation from the control values for both area and length were considered to change animal size and thus exclusion criteria were set to >3% change in area and > 2.5% change in length. If a given hit compound falls beyond both criteria it was considered as changed in body dimensions and excluded (10 hits in total). We performed a Pearson correlation to check for the validity of our exclusion criteria. For the 10 excluded drugs both area (R^2^=0.7), and length (R^2^=0.69) correlate with axon diameter changes. For the remaining 33 hits there was no correlation for area or length with diameter (R^2^=0.02 for area and R^2^=0.01 for length) (Figure 5).

### Validation of candidates

To validate hit compounds from the primary screen, the same compounds were ordered from different companies (Tocris, Cambridge Bioscience, Sigma, Santa Cruz, Cayman Chemicals). Embryos were again plated with 50 embryos per dish and were dechorinated if drug treatment started at 1 dpf. Depending on the price and amount of drug available, fish were treated in either 50 per dish or 96 wells with three fish per well. Clutch-specific DMSO-treated controls were included in every experiment and handled the same way as drug-treated fish. If the original LOPAC®^1280^ compound was still available, it was included and used as a positive control at its original dilution. Drugs were serially diluted according to the dilution used in the primary screen, ranging from 1, 2, 4, 10, 20, 50 to 100 µM. Treatment started either at 1 dpf or 3dpf with daily compound exchanges. Dilutions that resulted in unhealthy fish are not shown. Only healthy-appearing fish with inflated swim bladders were subsequently imaged using super-resolution FAST imaging at Zeiss LSM880 confocal microscope.

### Live in vivo imaging

Larvae were anaesthetized with 600 µM tricaine in E3 medium and immobilized in 1.5% low melting-point agarose on a coverslip glass dish and covered with tricaine/E3 medium to keep fish anaesthetized. 2-tiled z-stacks (with optimal z-step) were obtained using a Zeiss LSM880 microscope with Airyscan FAST in super-resolution mode, using a 40× objective lens (Zeiss Plan-Apochromat 40× water-dipping, NA = 1.0), and processed using the default Airyscan processing settings (Zen Black 2.3, Zeiss). All Mauthner images were taken from a lateral view of the spinal cord centered around somite 15-16. All lateral view images depict the anterior on the left and dorsal of the top, while dorsal view images depict the anterior on the top. Figure panels were prepared using Fiji (v1.51n) and Adobe Illustrator 2020 (24.0.2).

### Manual and script-based quantification of diameter

#### Manual Quantification for VAST-SDCM images

To quantify Mauthner axon diameter manually, z-stacks were loaded into Fiji/ImageJ (v1.51n). The look-up table (LUT) was set to “Thermal”, and images were thresholded using Min/Max. Only the top Mauthner axon (closest to the objective) was analysed to ensure consistency with the automated image analysis pipeline. A random grid with 500µm^2^ spacing was overlaid and every second square intersecting the top axon along its longitudinal axis was selected for measurement. Within each selected square, the Mauthner axon outline (identified by thermal LUT-highlighted turquoise pixels) was manually traced using polygon selection. A minimum of five regions distributed along the axon’s length were traced per sample. To calculate diameter, the area was divided by the corresponding width for each individual segment and the mean values across those 5-6 segments was used for analysis. A mean from those 5-6 diameter measurement was taken as the value. Because the manual measurements and the machine learning-based pipeline were trained and performed by different individuals, slight differences in axon outline definitions were expected (e.g., manual tracing based on thermal LUT-highlighted turquoise pixels versus training-based segmentation in the machine learning model). To account for this variability, axon diameter measurements were normalized by calculating the percentage change relative to the clutch-specific controls. Normalized diameter changes were then plotted as percentage differences relative to control.

#### Script based ImageJ Quantification for AiryScan FAST 880 confocal images

Axon diameter from AiryScan FAST confocal images were measured using custom-written ImageJ Macros and Fiji as described in detail in Bin et al^43^. Briefly, a “Split Axons Tool” was used to split the two adjacent Mauthner axons in whole larvae z-stack datasets, resulting in two separate maximum intensity projections for each Mauthner axon. Next, the “Axon Trace Tool” was used to trace the approximate mid-point of the axon along its length according to its intensity profile. Afterwards, the “Axon Calibre Tool” was used to measure the average axon diameter along the length of the selected Mauthner axon. The diameter was measured only for the axon located closest to the imaging objective. To account for clutch-specific variability we normalized axon diameter by calculating the change relative to DMSO-treated controls of the same clutch.

### Statistical analyses

Statistical analyses were performed using GraphPadPrism 10, as well as a combination of Python-based scripts and Excel-based module scripting to accommodate the large volume and complexity of screening data. p-Values are included in the figure legends and values for z-Score ranking and p-values for the hit compounds are included in Table 2. Significance was defined as p<0.05. Unless stated otherwise, n number represents values from independent fish and is included in the figure legends or the Supplementary Table 2. Only the one axon closest to the objective was quantified.

## Acknowledgements

We thank the University of Edinburgh BVS Zebrafish Facility and Zebrafish Imaging and Screening Facility for expert assistance. This work was supported by Wellcome Trust Senior Research Fellowships (214244/Z/18/Z) and a UKRI Frontier Research grant (EP/Z533890/1) to D.A.L.. M.A.E.V. is a recipient of a Walter-Benjamin Fellowship by the Deutsche Forschungsgemeinschaft (DFG grant 493410640 to M.A.E.V.) and a UKRI postdoctoral fellowship (EP/Y029577/1 to M.A.E.V.).

